# Emotional Content and Semantic Structure of Dialogues Are Associated With Interpersonal Neural Synchrony in the Prefrontal Cortex

**DOI:** 10.1101/2024.02.15.580458

**Authors:** Alessandro Carollo, Massimo Stella, Mengyu Lim, Andrea Bizzego, Gianluca Esposito

## Abstract

A fundamental characteristic of social exchanges is the synchronization of individuals’ behaviors, physiological responses, and neural activity. However, the association between how individuals communicate in terms of emotional content and expressed associative knowledge and interpersonal synchrony has been scarcely investigated so far. This study addresses this research gap by bridging recent advances in cognitive neuroscience data, affective computing, and cognitive data science frameworks. Using functional near-infrared spectroscopy (fNIRS) hyperscanning, prefrontal neural data were collected during social interactions involving 84 participants (i.e., 42 dyads) aged 18-35 years. Wavelet transform coherence was used to assess interpersonal neural synchrony between participants. We used manual transcription of dialogues and automated methods to codify transcriptions as emotional levels and syntactic/semantic networks. Our quantitative findings reveal higher than random expectations levels of interpersonal neural synchrony in the superior frontal gyrus (*q* = .038) and the bilateral middle frontal gyri (*q* ***<*** .001, *q* ***<*** .001). Linear mixed models based on dialogues’ emotional content only significantly predicted interpersonal neural synchrony across the prefrontal cortex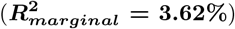 . Conversely, models relying on syntactic/semantic features were more effective at the local level, for predicting brain synchrony in the right middle frontal gyrus 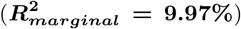 Generally, models based on the emotional content of dialogues were not effective when limited to data from one region of interest at a time, whereas models based on syntactic/semantic features show the opposite trend, losing predictive power when incorporating data from all regions of interest. Moreover, we found an interplay between emotions and associative knowledge in predicting brain synchrony, providing quantitative support to the major role played by these linguistic components in social interactions and in prefrontal processes. Our study identifies a mind-brain duality in emotions and associative knowledge reflecting neural synchrony levels, opening new ways for investigating human interactions.

## 1 Introduction

Social interactions serve as catalysts for human development and well-being. As in many other social species, human newborns need a significant other to regulate their physiological states and guarantee their survival [1]. Ever since childhood, individuals engage in social exchanges transmitting cognitive, social, and emotional competencies or skills [2, 3]. Newborns are inherently prepared for this task. Extensive research in cognitive neuroscience has shown that evolution has equipped newborns with a series of mechanisms that enable them to pay attention and adequately respond to the social world around them [4]. Importantly, social interactions remain fundamental in adulthood, with the quality of one’s social exchanges being a predictive factor of both the susceptibility to psychiatric disorders and the mortality risk [5, 6].

Recognizing the paramount importance of social interactions in people’s lives, numerous studies have investigated the characteristics of adaptive social exchanges (e.g., a mother talking to their child or teachers pointing at some writing on a blackboard) [7]. In particular, the mutual attunement of behavioral and physiological signals between interactive partners, known as *bio-behavioral synchrony*, has emerged as a fundamental mechanism through which social interactions influence individuals’ development and well-being [8–10]. Traditional research on synchrony has predominantly explored behaviors, hormonal fluctuations, and physiological responses, reporting attunement over these multiple levels during social exchanges [11]. Recently, the introduction of the hyperscanning approach in cognitive neuroscience has expanded the *bio-behavioral synchrony* framework to go beyond behavioral and physiological data, including the analysis of the neural level [12]. Based on the simultaneous recording of brain activity in two (or even more) individuals, the hyperscanning approach allows the investigation of the neural underpinnings of dyadic (or group-level) social interactions [13]. Initial hyperscanning studies used traditional neuroimaging techniques (e.g., electroencephalography (EEG), functional magnetic resonance imaging (fMRI)) to assess instances of interpersonal neural synchrony (e.g., [13, 14]). In these studies, participants interacted in highly controlled experimental situations because of the characteristics of the adopted neuroimaging instrument. In fact, both EEG and fMRI require participants to minimize their body movements in order to collect usable neural data. Moreover, fMRI requires that participants lay supine within a scanner. Altogether, these features did not allow participants to interact as they would normally do in daily social exchanges. More recently, thanks to the advent of functional near-infrared spectroscopy (fNIRS), hyperscanning research has shown promise for the study of social interactions in real-life scenarios (e.g., [15–17]). Recent fNIRS hyper-scanning studies have demonstrated that interpersonal neural synchrony can serve as a marker of the quality of relational social exchanges [18, 19].

In recent hyperscanning studies, tasks replicating naturalistic interactions, such as mother-child free play, have become increasingly prevalent (e.g., [20, 21]). Despite this trend, the exploration of the affective component, which constitutes the core of real-life social interactions, in the context of interpersonal neural synchrony remains largely unexplored [22–24]. The affective dimension of social exchanges plays a pivotal role in various aspects, including the establishment of social cohesion and the determination of one’s inclination toward engaging in prosocial behaviors [25, 26]. Notably, some seminal studies have investigated the association between emotions and interpersonal neural synchrony. These studies adopted a framework based on the circumplex model of affect in neuroscience [27], which maps human emotions along two dimensions: *(i)* pleasantness/unpleasantness (valence) and *(ii)* activation/inhibition (arousal). The works by [20], by [28], and [29] showed that higher emotional valence of stimuli or interactions is associated with higher interpersonal neural synchrony. In other words, the presence of emotions eliciting more extreme pleasantness or unpleasantness corresponded to higher coherence between individuals in terms of their respective brain signals. However, the direction of the association between the valence of emotions and interpersonal neural synchrony is unclear. Some studies report higher interpersonal neural synchrony for negatively-valenced emotional content (e.g., [28]), while others reported the same effect for positively-valenced emotional content (e.g., [29, 30]). Furthermore, [28] observed that arousal also correlates positively with interpersonal neural synchrony. In their recent theoretical framework, [31] identify the ability to elicit emotional responses as one of the stimulus characteristics that enhance interpersonal neural synchrony. According to the authors, emotional stimuli may achieve this through different mechanisms. For example, they might exert an indirect effect by altering the perception of time and its passage. Alternatively, emotional stimuli could directly foster synchrony by enhancing overall stimulus processing and aligning internal rhythms with external ones. Consequently, the authors propose that emotional stimuli – particularly positive ones – play a preparatory role in facilitating the representation of another person’s thoughts, feelings, and behaviors, including their temporal dynamics. However, these and similar findings were relative to individuals self-reporting their emotional states or to emotional states induced by an emotional stimulus, whereas future research should concentrate on detecting emotions from potential language-based interactions at play while measuring neural synchrony.

A key channel through which emotions are conveyed between individuals is language [32]. Language represents an evolutionary achievement enabling the sharing of communicative intentions between individuals. These intentions encode ideas and emotions either in written (i.e., text) or verbal form (i.e., speech). Speech is a complex system [33, 34], where spoken words can be assembled to denote more elaborate ideas, with distinctive non-cognitive features, like emotions [35]. Speech does not depend only on its “bag” of words [36] (i.e., words in a speech or text) but also on the structure of word-word relationships [35, 37] (e.g., on the syntactic links mixing words to convey efficiently meanings and emotional states) [38]. How people communicate might be more or less effective depending on the specific structure of conceptual relationships and emotional eliciting employed in dialogues. For instance, when conversing, people can superficially explore different topics or, conversely, they can dive into a deep discussion of one topic of interest [34, 39]. Along the theory of basic human emotions developed by [40] and mapping a variety of nuanced emotions as combinations of 8 basic emotional states, cognitive science [38, 41, 42] and neuroscience [8, 43] pose converging evidence that three key features of language might influence the quality of the social exchange as well as the level of interpersonal neural synchrony: *(i)* the emotional content (e.g., a portion of language eliciting emotions like anger or trust); *(ii)* the syntactic and semantic organization of ideas/concepts capturing the structure of associative knowledge as coded in the theory of the mental lexicon [44] (e.g., synonyms being used to specify similar ideas); *(iii)* frequency effects (e.g., turn-taking frequency of quips). In terms of frequency effects, some recent studies have already shown that the rhythm of the interaction, as measured in terms of turn-taking frequency, relates to interpersonal neural synchrony in mother-child dyads with verbal and even pre-verbal children [45, 46]. However, the effect of the emotional content and syntactic/semantic structure remains scarcely investigated.

This paper aims to fill this research gap in the relevant literature. To do so, we introduce a quantitative framework investigating the two dimensions of dialogues’ emotional content and syntactic/semantic structure for investigating their association with interpersonal neural synchrony during naturalistic social exchanges. Our study, thus, bridges innovative cognitive neuroscience data [15, 47] with affective computing and cognitive data science frameworks [35], integrating mind and brain data [48, 49]. In particular, we leverage the recent modeling framework of *Textual Forma Mentis Networks* (TFMNs) for representing conceptual associations encoded in text and retrieved via artificial intelligence (AI) and psychologically validated data [37, 50]. TFMNs can encode associated mindsets (e.g., ways of associating concepts together).

We hypothesize that:

1. The emotional content of dialogues is significantly associated with interpersonal neural synchrony. Specifically, we hypothesize that higher expression of emotions would related to higher interpersonal neural synchrony in dialogues;
2. The structure of syntactic/semantic associations between concepts in dialogues is significantly associated with interpersonal neural synchrony.

To explore the interplay between these two dimensions, we use recently available data from a fNIRS hyperscanning study, under three conditions (i.e., natural conversation, role-play, and role reversal) based on dialogues among participants [15, 47, 51]. The effects of these conditions on interpersonal neural synchrony have already been explored in previous studies [15, 51], and the dataset was selected for its richness in dialogue interactions. For this reason, in the current work, we do not make predictions related to the conditions and do not test their effect on synchrony.

## 2 Results

Before investigating the association between emotional content and syntactic/semantic structure of dialogues and interpersonal neural synchrony, a preliminary analysis of synchrony across real (face-to-face conversationalists) and surrogate (randomly-paired and non-interacting) dyads for each region of interest (i.e., anterior prefrontal cortex, superior frontal gyrus, left middle frontal gyrus, right middle frontal gyrus, left inferior frontal gyrus, and right inferior frontal gyrus) was conducted. In this preliminary analysis, the goal was to identify the regions of interest that showed above-chance levels of synchrony scores in order to use them in subsequent analyses.

To investigate the association between emotional content of dialogues and inter-personal neural synchrony, we conducted a series of linear mixed models. In particular, we conducted one linear mixed model for data from individual regions of interest, with WTC scores as the dependent variable, the eight basic emotions encoded in EmoAtlas [52] as fixed predictors, and the experimental condition and dyad ID as random effects. With these models, we were able to disentangle the relationship between specific emotions and synchrony in specific regions of interest. Another linear mixed model was, then, computed on data from all the regions of interest by adding the region of interest as random effect. We used the same approach with the syntactic/semantic structure of dialogues. Finally, we combined information regarding both the emotional content and the syntactic/semantic structure of dialogues to predict the interpersonal brain synchrony in the whole prefrontal cortex.

### 2.1 Interpersonal Neural Synchrony Across True and Surrogate Dyads

Before investigating the relationship between the emotional content and the syntactic/semantic structure of dialogues, we conducted a preliminary analysis of synchrony scores. This allowed us to identify and select only the regions of interest that exhibited above-chance levels of synchrony for use in subsequent analyses. To achieve this, we conducted six linear mixed models – one for each region of interest – using WTC scores as the dependent variable. We performed Benjamini-Hochberg false discovery rate correction to control for Type I errors due to multiple tests. The fixed predictor was dyad type (true *versus* surrogate), while the experimental condition and dyad ID were included as random effects.

Statistically significant differences emerged between real and surrogate dyads in the WTC in their superior frontal gyrus (*β* = 0.010, *Standard Error (SE)* = 0.005, *t*(207) = 2.201, *p* = .029, *q* = .038), left middle frontal gyrus (*β* = 0.018, *SE* = 0.004, *t*(203.50) = 4.196, *p <* .001, *q <* .001), and right middle frontal gyrus (*β* = 0.017, *SE* = 0.005, *t*(202.20) = 3.468, *p <* .001, *q <* .001); see Figure 1. No statistically significant difference (*q >* .05) emerged in the WTC computed in the anterior prefrontal cortex, left inferior frontal gyrus, and right inferior frontal gyrus. For this reason, the subsequent statistical analyses will exclude WTC from these regions of interest that did not show significant differences between true and surrogate dyads.

**Fig 1.**
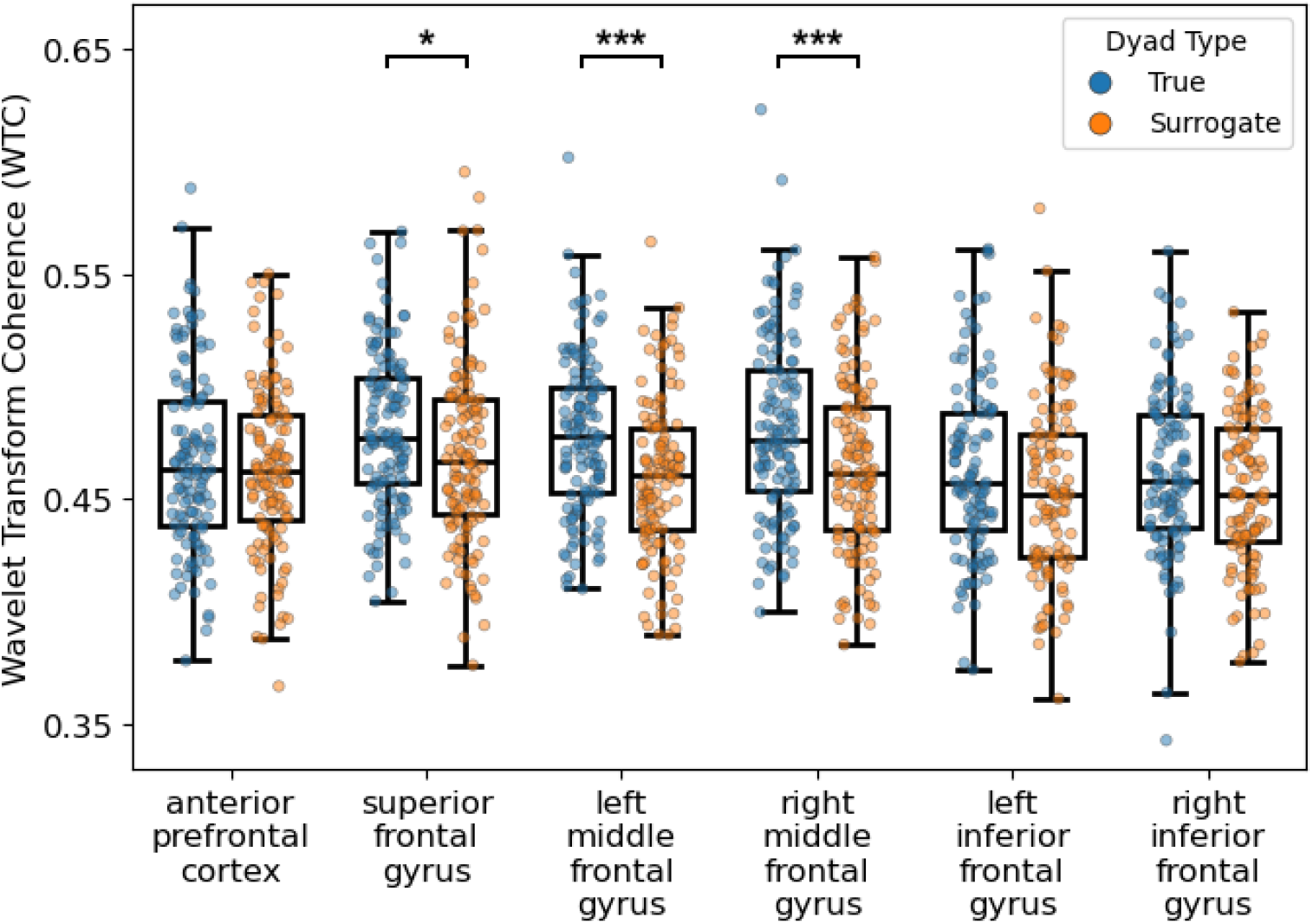
Wavelet Transform Cohesrence (WTC) values across regions of interest in the prefrontal cortex. For each region of interest, values of WTC in true (in blue) and surrogate (in orange) dyads are provided. (* *q <* .05; *** *q <* .001).

### 2.2 Emotional Content of Dialogues and Interpersonal Neural Synchrony

In this analysis, we used emotional *z* -scores provided by EmoAtlas (see Section 5.7) as proxies of the emotional content of dialogues. Figure 2 illustrates the distribution of aggregated emotional *z* -scores in the dataset. As shown in the Figure, the most frequently expressed emotions were anticipation and joy, whereas anger, disgust, and fear were the least expressed.

**Fig 2.**
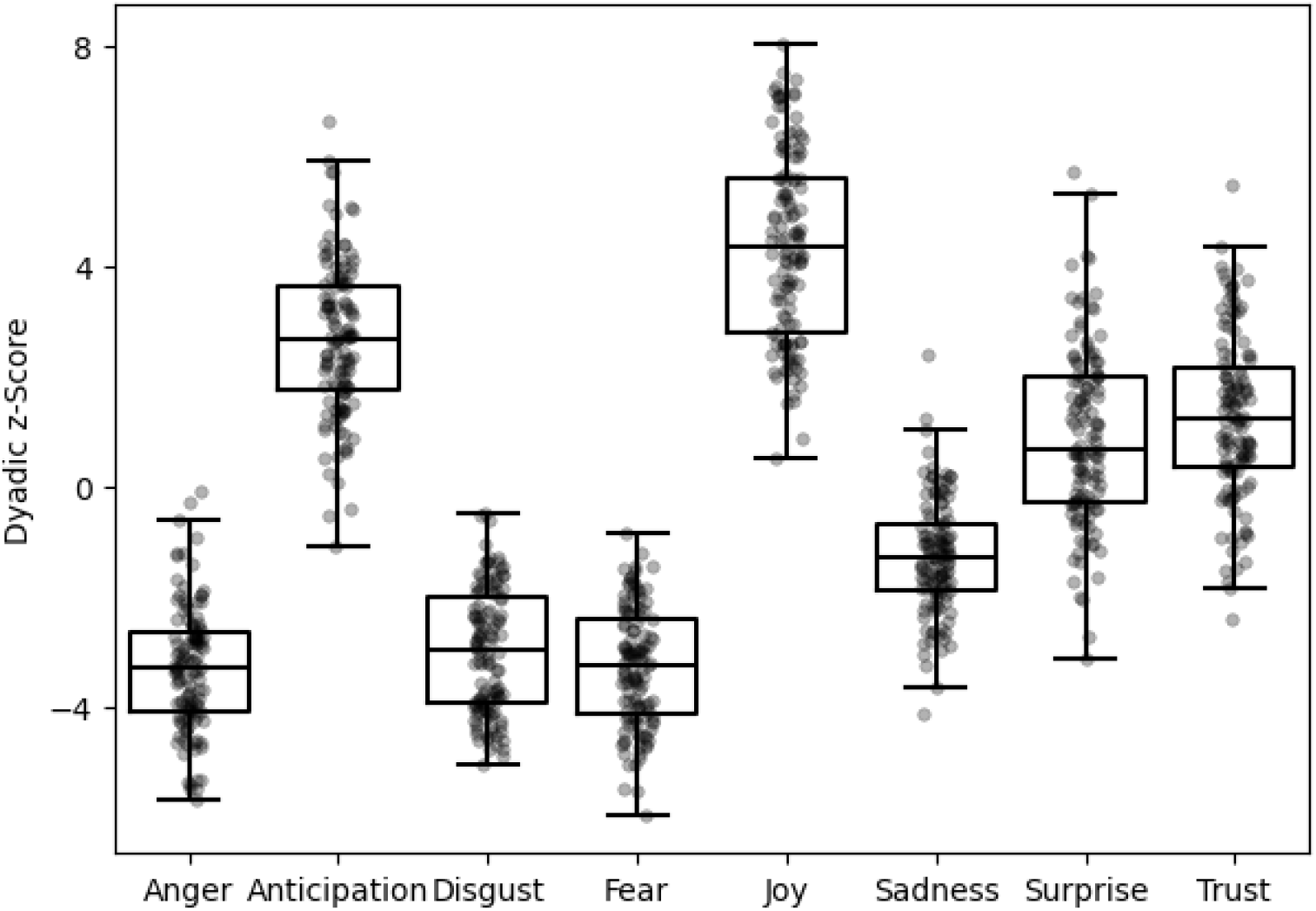
Distribution of aggregated emotional *z* -scorses representing the emotional content of the transcribed dialogues.

#### 2.2.1 Analysis on Individual Regions of Interest

Three linear mixed models were conducted to analyze data from three regions of interest that exhibited above-chance levels of synchrony: the superior frontal gyrus, the left middle frontal gyrus, and the right middle frontal gyrus. In each model, WTC scores were used as the dependent variable, emotional *z* -scores as fixed effects, and the experimental condition and dyad ID as random effects.

In the three models, emotional *z* -scores were not significant predictors of interpersonal neural synchrony (*q >* .05); see Table 1 for a summary of the results).

**Table 1.** Summary of fixed effects related to the emotion *z* -scores in the linear mixed models conducted to predict interpersonal neural synchrony in individual regions of interest (i.e., superior frontal gyrus, left middle frontal gyrus, right middle frontal gyrus). For each emotion, we reported the estimate, the standard error, the *t* -value, the *p*-value, and the *q* -value. Statistically significant *q* -values are noted in bold.

| Predictor | Estimate ($\beta$) | Standard Error | $t$-value | $p$-value | $q$-value |
| --- | --- | --- | --- | --- | --- |
| **Interpersonal Neural Synchrony on the Superior Frontal Gyrus** |  |  |  |  |  |
| Anger | -0.0037 | 0.005 | -0.796 | .428 | .730 |
| Anticipation | -0.0026 | 0.004 | -0.668 | .505 | .744 |
| Disgust | -0.0002 | 0.004 | -0.039 | .969 | .971 |
| Fear | 0.0062 | 0.004 | 1.551 | .124 | .372 |
| Joy | 0.0041 | 0.004 | 1.148 | .254 | .684 |
| Sadness | 0.0001 | 0.004 | 0.037 | .971 | .971 |
| Surprise | 0.0024 | 0.004 | 0.598 | .551 | .744 |
| Trust | 0.0064 | 0.003 | 1.852 | .067 | .300 |
| **Interpersonal Neural Synchrony on the Left Middle Frontal Gyrus** |  |  |  |  |  |
| Anger | -0.0092 | 0.005 | -1.926 | .057 | .300 |
| Anticipation | -0.0080 | 0.004 | -2.002 | .048 | .300 |
| Disgust | 0.0075 | 0.004 | 1.675 | .097 | .327 |
| Fear | -0.0010 | 0.004 | -0.241 | .810 | .875 |
| Joy | 0.0027 | 0.004 | 0.751 | .454 | .730 |
| Sadness | 0.0030 | 0.004 | 0.759 | .450 | .730 |
| Surprise | 0.0038 | 0.004 | 0.962 | .338 | .730 |
| Trust | 0.0033 | 0.004 | 0.935 | .352 | .730 |
| **Interpersonal Neural Synchrony on the Right Middle Frontal Gyrus** |  |  |  |  |  |
| Anger | -0.0042 | 0.006 | -0.742 | .460 | .730 |
| Anticipation | -0.0082 | 0.005 | -1.677 | .096 | .327 |
| Disgust | 0.0024 | 0.005 | 0.453 | .651 | .837 |
| Fear | -0.0014 | 0.005 | -0.289 | .773 | .870 |
| Joy | -0.0014 | 0.004 | -0.316 | .753 | .870 |
| Sadness | 0.0028 | 0.005 | 0.604 | .547 | .744 |
| Surprise | 0.0017 | 0.005 | 0.358 | .721 | .870 |
| Trust | -0.0045 | 0.004 | -1.058 | .293 | .718 |

#### 2.2.2 Analysis on the Whole Prefrontal Cortex

When analyzing all scores from the prefrontal cortex, we can either put all regions of interest together or perform the analysis on each region of interest separately.

To explore the relationship between prefrontal interpersonal neural synchrony and the emotional content of dialogues, we conducted a linear mixed model. In this model, WTC scores served as the dependent variable, emotional *z* -scores were included as fixed effects, and region of interest, experimental condition, and dyad ID were included as random effects. Results indicated that anticipation was a significant predictor of interpersonal neural synchrony (*β* = −0.007, *SE* = 0.003, *t*(247.98) = -2.695, *p* = .008, *q* = .025; see Table 2 for an overview of the results). The *AIC* of the model equaled to -1223.85 and the 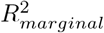 was equal to 3.62%.

**Table 2.**
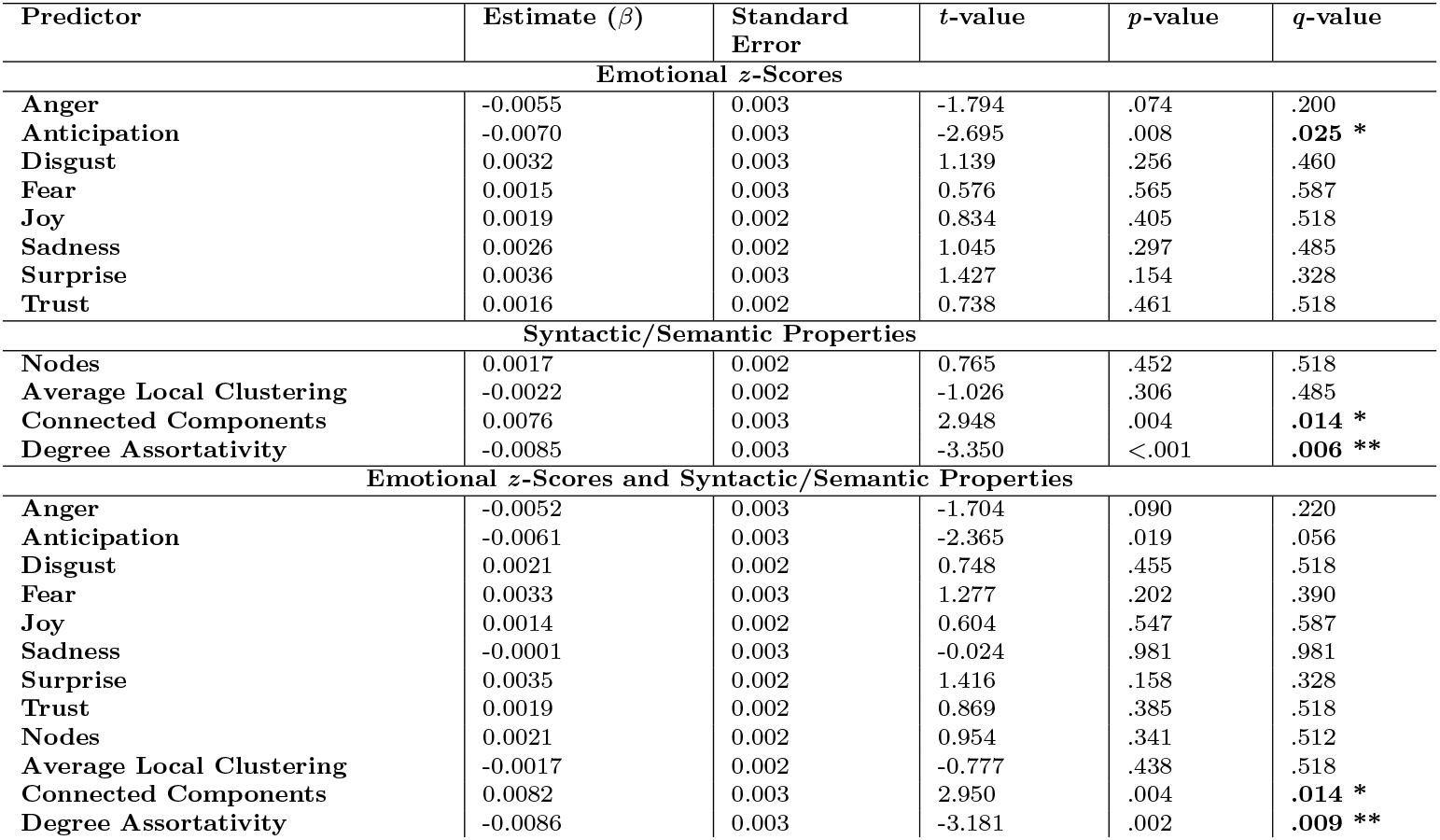
Summary of fixed effects in the linear mixed models conducted to predict interpersonal neural synchrony using data from all regions of interest (i.e., superior frontal gyrus, left middle frontal gyrus, right middle frontal gyrus). For each predictor, we reported the estimate, the standard error, the *t* -value, the *p*-value, and the *q* -value. Statistically significant *q* -values are noted in bold. (* *q <* .05; ** *q <* .01).

### 2.3 Syntactic/Semantic Structure of Dialogues and Interpersonal Neural Synchrony

In this analysis, we employ TFMNs (see Section 5.8) as proxies of the syntactic/semantic associations between concepts encoded within communicative intentions.

#### 2.3.1 Analysis on Individual Regions of Interest

Three linear mixed models were conducted to analyze data from three regions of interest that exhibited above-chance levels of synchrony: the superior frontal gyrus, the left middle frontal gyrus, and the right middle frontal gyrus. In each model, WTC scores were used as the dependent variable, syntactic/semantic metrics as fixed effects, and the experimental condition and dyad ID as random effects.

One out of the three models showed a significant effect of syntactic/semantic properties of dialogues on interpersonal neural synchrony (see Table 3 for a summary of the results). Specifically, the model on data from the right middle frontal gyrus showed a significant effect of the degree assortativity (*β* = −0.015, *SE* = 0.006, *t*(106.25) = -3.361, *p* = .001, *q* = .004) on WTC scores. The *AIC* of the model equaled to -365.83 and the 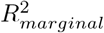 was equal to 9.97%. In contrast, syntactic/semantic metrics were not significant predictors of interpersonal neural synchrony in the models on data from the superior frontal gyrus and from the left middle frontal gyrus (*q >* .05).

**Table 3.** Summary of fixed effects related to the syntactic/semantic properties of dialogues in the linear mixed models conducted to predict interpersonal neural synchrony in individual regions of interest (i.e., superior frontal gyrus, left middle frontal gyrus, right middle frontal gyrus). For each syntactic/semantic metric, we reported the estimate, the standard error, the *t* -value, the *p*-value, and the *q* -value. Statistically significant *q* -values are noted in bold. (** *q <* .01).

| Predictor | Estimate ($\beta$) | Standard Error | $t$-value | $p$-value | $q$-value |
| --- | --- | --- | --- | --- | --- |
| **Interpersonal Neural Synchrony on the Superior Frontal Gyrus** |  |  |  |  |  |
| Nodes | -0.0012 | 0.003 | -0.370 | .712 | .801 |
| Average Local Clustering | 0.0003 | 0.003 | 0.094 | .925 | .925 |
| Connected Components | 0.0027 | 0.004 | 0.692 | .491 | .669 |
| Degree Assortativity | 0.0013 | 0.004 | 0.322 | .748 | .801 |
| **Interpersonal Neural Synchrony on the Left Middle Frontal Gyrus** |  |  |  |  |  |
| Nodes | 0.0028 | 0.003 | 0.821 | .418 | .627 |
| Average Local Clustering | -0.0034 | 0.003 | -1.007 | .316 | .527 |
| Connected Components | 0.0073 | 0.004 | 1.804 | .074 | .158 |
| Degree Assortativity | -0.0091 | 0.004 | -2.256 | .026 | .065 |
| **Interpersonal Neural Synchrony on the Right Middle Frontal Gyrus** |  |  |  |  |  |
| Nodes | 0.0023 | 0.004 | 0.600 | .550 | .688 |
| Average Local Clustering | -0.0041 | 0.004 | -1.104 | .272 | .510 |
| Connected Components | 0.0111 | 0.005 | 2.433 | .017 | .050 |
| Degree Assortativity | -0.0152 | 0.006 | -3.361 | .001 | **.004\*\*** |

Comparing all the above results, we notice that the emotional content of conversations does not act as a significant predictor of interpersonal neural synchrony in individual regions of the prefrontal cortex. Instead, syntactic/semantic properties are significant predictors of brain synchrony already at the local level (i.e. in the right regions of the prefrontal cortex). This distinctiveness calls for additional inquiry within the Discussion.

#### 2.3.2 Analysis on the Whole Prefrontal Cortex

To investigate the relationship between prefrontal interpersonal neural synchrony and the syntactic/semantic properties of dialogues, we conducted a linear mixed model. In this model, WTC scores were used as the dependent variable, syntactic/semantic metrics were included as fixed effects, and region of interest, experimental condition, and dyad ID were included as random effects. We observed a significant effect of the number of connected components (*β* = 0.008, *SE* = 0.003, *t*(218.30) = 2.948, *p* = .004, *q* = .014) and degree assortativity (*β* = −0.009, *SE* = 0.003, *t*(247.88) = -3.350, *p <* .001, *q* = .006) on neural synchrony (see Table 2 for an overview of the results).The *AIC* of the model equaled to -1275.416 and the 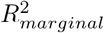 was equal to 4.44%.

Two key observations can be drawn from these results. First, models based on syntactic/semantic metrics tend to lose predictive power when applied to all regions of interest, compared to when they are focused on specific portions of the prefrontal cortex. Second, comprehensive models utilizing syntactic/semantic information to predict synchrony across the entire prefrontal cortex exhibit slightly greater predictive power than those based solely on emotional information.

### 2.4 Emotional Content, Syntactic/Semantic Structure of Dialogues, and Interpersonal Neural Synchrony

Finally, we combined information regarding both the emotional content and the syntactic/semantic structure of dialogues to predict the interpersonal brain synchrony in the whole prefrontal cortex. The analysis revealed a significant effect of the number of connected components (*β* = 0.008, *SE* = 0.003, *t*(205.40) = 2.950, *p* = .004, *q* = 014), and degree assortativity (*β* = − 0.009, *SE* = 0.003, *t*(245.40) = -3.181, *p* = .002, *q* = .009) on WTC scores (see Table 2). The *AIC* of the model equaled to -1187.726 and 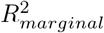 the was equal to 7.60%.

We observe that, compared to models that rely solely on emotional information or syntactic/semantic information as primary predictors of synchrony in the prefrontal cortex, the combined model that integrates both types of information demonstrates superior performance and greater predictive power.

## 3 Discussion

The current study investigated how emotions and associative knowledge in dialogues relates to interpersonal neural synchrony during a hyperscanning experiment. Prefrontal neural activity data were measured with fNIRS. Pre-recorded dialogues [15, 47] among social dyads were manually transcribed and investigated. Automated methods of emotional content detection and syntactic/semantic structure mapping were adopted [37, 52]. By using this mind-brain approach, we observe that the emotional content and the syntactic/semantic structure of dialogues significantly associates with interpersonal neural synchrony across regions of the prefrontal cortex: the superior frontal gyrus and the bilateral middle frontal gyri.

Firstly, we observe that, across the whole prefrontal cortex, these regions of interest show a significant difference in interpersonal neural synchrony between real *versus* surrogate dyads. Our findings agree with past results observed by the pioneering hyperscanning study conducted by [53]. More in detail, Cui and colleagues found that the superior frontal cortex shows increased interpersonal neural synchrony in cooperative tasks different from the experimental conditions tested here (i.e., computer-based cooperation game). As for the authors, these portions of the prefrontal cortex could have a role in controlling processes related to coordination with others within the theory of mind [54]. Conversely, brain synchrony in the anterior prefrontal cortex and the bilateral inferior frontal gyri does not appear to differ between true and surrogate dyads across the experiment. This result contrasts with previous hyperscanning studies that found above-chance levels of brain synchrony in these regions (e.g., [45, 55]). These differing patterns might be explained by the specific conditions of the current experiment, where participants had to interact while playing the role of another persona. In fact, the prefrontal cortex is involved in situations of identity conflict and faking [47, 56–58], with the inferior frontal gyrus being more involved in identity concealment, and the medial regions of the prefrontal cortex being more involved during identity faking [57, 58]. Among the two conditions, identity faking is the one that resembles the tasks of the current study.

Moreover, in interacting individuals, we find that the synchronization between the superior frontal gyrus and the bilateral middle frontal gyri is associated with both the emotional content and the syntactic/semantic structure of dialogues. This finding seems in contrast to the results of the EEG hyperscanning study by [59], in which the authors observed that interpersonal brain synchrony did not vary based on emotional, reminiscent, and practical aspects of conversations. However, Kinreich and colleagues considered these aspects as being binary variables (i.e., either present or absent according to human raters). Our analysis adopts a different human-centered AI approach, going beyond binary characterizations and reconstructing the complex network structure of emotions and associative knowledge expressed in dialogues. This finer level of resolution, enabled by more recent natural language processing techniques, can explain such a difference. Speech in dialogues represents a complex system [35] where naturalistic social interactions convey various pieces of information beyond the content alone. For example, dialogues can be characterized by the emotions that they convey [40] or by the syntactic/semantic associations between ideas expressed by the speaker [34].

In the present study, the emotional content did not significantly relate to the activity of a single individual region of interest. However, emotions in dialogues associates with interpersonal neural synchrony in the superior frontal gyrus and the left middle frontal gyrus when these regions are considered together. Both these regions are known in the literature for being involved in emotional processing and regulation strategies [60]. The superior frontal gyrus tends to show higher activation when the person up-or down-regulates emotional states [61]. The rostral part of the middle frontal gyrus – which is a part we explored with fNIRS – is involved in the meaning-making of emotional stimuli that depend on salience and relevance for the self [62, 63]. In other words, this brain region assigns a meaning to stimuli based on self-related implications. Indeed, the rostral middle frontal gyrus also shows prolonged activity in response to emotional events [64]. To testify for their role in emotional processing, both the left middle frontal gyrus and superior frontal gyrus show higher spontaneous activity in individuals with affective disorders (e.g., major depressive disorder) [65] and when people assess the emotional expression of music [66].

Dialogues’ syntactic/semantic structure highlights different patterns compared to emotional content. Higher interpersonal neural synchrony in the right middle frontal gyrus is associated with negative values of degree assortativity [67]. Therefore, couples with higher interpersonal neural synchrony in the right middle frontal gyrus tended to link highly connected words (e.g., frequent or general terms) with less connected ones (e.g., more infrequent or specific terms). This trend might suggest a relationship between language entropy and interpersonal neural synchrony in the right middle frontal gyrus. Regarding the observed results, previous studies have shown that the middle frontal gyrus is involved in syntactic/semantic processes. This brain region shows enhanced activity for semantically distant or unrelated words when compared to activity elicited by semantically similar or related words [68–70]. More specifically, the activity in the right portion of the middle frontal gyrus is reportedly modulated by categorical relationships among words [71–73]. Hence, our findings agree with past results indicating that the middle frontal gyrus should intervene when processing words with different syntactic and semantic connectivity.

When predicting interpersonal neural synchrony across the prefrontal cortex, models based on emotional content and syntactic/semantic properties of dialogues outperform their counterparts (e.g., models using emotional information or syntactic/semantic network features alone). Moreover, models based on syntactic/semantic network features lose predictive power as compared to when they are used with data from individual regions of interest. This pattern supports the notion for which the prefrontal cortex is involved in both emotional and syntactic processes [74, 75], with each source of information contributing in explaining the brain activity (and synchronization) in this region. Moreover, our findings relate to the difference in global and local processing of emotional and syntactic/semantic aspects in the prefrontal cortex during social interactions. On one hand, the emotional content of conversations appears to be a predictor of brain synchrony across the whole prefrontal cortex and only marginally of synchrony in specific subregions. Emotional information possibly requires larger samples to demonstrate its predictive power at the local level of analysis. Conversely, syntactic/semantic features of dialogues emerge as better predictors of brain synchrony at the local level, specifically in the right middle frontal gyrus. When introducing scores of brain synchrony from other regions of the prefrontal cortex, models based on syntactic/semantic properties tend to show lower performance, probably due to the introduction of noise.

### 3.1 Limitations and Future Research

In discussing the results of the study, it is important to acknowledge certain aspects that should be considered as potential limitations and serve as a foundation for future research.

Firstly, our investigation utilized fNIRS hyperscanning to assess neural activity in prefrontal cortex regions. While the prefrontal cortex is recognized for its involvement in social processes [76], it is now known that other brain regions (e.g., temporo-parietal junction) exhibit interpersonal neural synchrony [19], which might be influenced by factors such as emotional content and the syntactic/semantic structure of dialogues, especially in natural conversations. Moving forward, potential lines of research could expand upon the current findings by exploring other brain regions exhibiting interpersonal neural synchrony.

Secondly, our study employed the emotional content of dialogues as a predictor of interpersonal neural synchrony. However, emotions are not solely conveyed through speech content [40]. Rather, an individual’s emotional state may emerge through non-verbal cues or subtle indicators, such as facial expressions or tone of voice. Future works might capitalize on recent developments in cognitive data science to assess automatically visual and auditory channels through which emotions are conveyed among humans [77].

Thirdly, our study focuses only on one socio-cultural set of conditions (e.g., close friends within the Italian population). This limitation was mostly due to technical issues with sampling individuals through hyperscanning. Additionally, we did not further investigate the degree or depth of the friendship within the dyads, which could potentially influence interpersonal neural synchrony. Future research could involve different types of dyads (e.g., romantic partners, strangers, parent-child relationships) to see how our current results generalize across different types of social bonds and potentially different levels of empathy.

Fourthly, another limitation of the study is that the experimental design [15, 51] focused on positive scenarios. However, thanks to the model, we are able not only to detect the presence (*z* -score *>* 1.96) or absence ( |*z* − *score* | *<* 1.96) of a specific emotion but we can also determine whether the participants actively avoid using jargon relative to a given emotional state (*z* -score *<*− 1.96). Hence, even though the design did not focus on negative emotional states, our model was still able to discriminate between dyads of participants actively avoiding negative emotions or not. Future research could design other experimental conditions eliciting different or negative emotions as coded in emotional theories like [40]’s or the circumplex model of affect [27].

Taking these limitations into account, the current work represents a pioneering and human-centered AI exploration into understanding the role of emotions and associated knowledge in predicting interpersonal neural synchrony.

## 4 Conclusion

The current fNIRS hyperscanning study has provided valuable insights into bridging neuroscientific, affective, and cognitive aspects of interpersonal synchrony building on past mind-brain approaches [15, 47–49, 51]. More in detail, we quantified how the emotional content and syntactic/semantic associations within dialogues relate to interpersonal neural synchrony in the prefrontal cortex. Specifically, the emotional content of conversations tended to show a consistent association with brain synchrony throughout the whole prefrontal cortex, losing its predictive power when analyzing data from smaller regions of interest. Conversely, associative knowledge structure [35] was found to have a stronger association with interpersonal neural synchrony in the right middle frontal gyrus, losing part of its predictive power when incorporating data from all the regions of the prefrontal cortex. Notably, when we focused on the entire prefrontal cortex, the best predictive model was the one that integrated both emotional content and semantic/syntactic information. This result highlights the combined relevance of affective and cognitive components in human interactions and in the activity of the prefrontal cortex. Our transdisciplinary data-informed approach corroborates the significance of the affective component in real-life social interactions, within the context of the *bio-behavioral synchrony* framework [10].

## 5 Methods

### 5.1 Study Design

All data for the study were obtained from a cross-cultural investigation testing the neural underpinnings of role-play [15, 47, 51], which is a clinical technique used to alleviate psychopathological symptoms. Data were collected with a hyperscanning approach based on fNIRS. Participants were asked to interact freely for 5 minutes in three different conditions: natural conversation, role-play, and role reversal. The same experiment was conducted in Italy and Singapore, but for the current study, we used only the data from the Italian cohort. We chose to focus only on the Italian data due to the linguistic variability present in the Singaporean cohort. Participants in Singapore interacted in multiple languages, often incorporating Singlish, a creole that is not supported by the model we used to extract dialogue features. The high linguistic variability in the Singaporean cohort could have introduced confounding factors, potentially influencing the results and increasing the risk of false discoveries driven by language differences rather than the intrinsic properties of the dialogues. All the interactions were recorded and manually transcribed. To identify dialogues’ emotional content and syntactic/semantic structure, we used automated methods such as EmoAtlas and TFMNs. Ultimately, emotional *z* -scores and the main properties of the syntactic/semantic network were used to predict interpersonal neural synchrony across different sub-regions of the prefrontal cortex of the brain. Data collections were approved by the University of Trento (2022-059) and by Nanyang Technological University (NTU-IRB-2021-03-013). The conduction of the experiment followed the guidelines provided by the Declaration of Helsinki. Informed consent was obtained from all participants.

### 5.2 Participants

As mentioned above, for the current study, we used only the data from the cohort in Italy (*N* = 84 participants, i.e., 42 dyads; age range = 18-35 years old). Participants were recruited via convenience and snowball sampling from social media sites. All the recruited dyads consisted of friends with an existing peer relationship. Participants reported no history of known and/or diagnosed health or neurological conditions, particularly those that could alter the oxygen-binding capacity of the blood. Additionally, none of the participants were using any medications, especially those with the potential to alter blood flow or blood pressure.

### 5.3 Experimental Protocol

The experimental procedure consisted of four separate phases, in which neural data were obtained employing fNIRS [15, 47, 51]. Firstly, the experiment began with two minutes of resting state to record the neural activity at rest. During the resting state, participants were asked to sit silently in front of each other and not to move their limbs as much as possible (see Figure 3A for the experimental setup). The subsequent phases were: (2nd) natural conversation, (3rd) role-play, and (4th) role reversal. Each of these phases lasted for five minutes. In the natural conversation condition, participants could interact as they would normally do in a daily conversation. In the role-play phase, participants were asked to interact by pretending to be another pair of mutual friends among their peer group. In the role reversal phase, participants were asked to interact by exchanging their roles (i.e., participant A pretending to be participant B, and *vice versa*). To provide a common framework for dialogues, a shared prompt was used across all interactive conditions. In this prompt, participants were instructed to pretend that they spotted each other at the shopping mall while trying to buy each other presents. To reduce primacy effects, the order of natural conversation, role-play, and role reversal was randomized. For all conditions, we truncated all data from the first and fifth minutes of interactions, as the beginnings and endings of conversations are structurally different from the main interaction [47, 78]. The resting phase was not used in the current analyses, as our focus was on conditions involving active dialogue to investigate the association between emotional content, syntactic/semantic properties of conversations, and interpersonal neural synchrony. Moreover, the effect of the interactive conditions on interpersonal neural synchrony has already been explored in previous studies [15, 51]. For this reason, we did not further test their effect on synchrony, as this would go beyond the scope of the current work, and controlled for the experimental conditions in the statistical models by using them as random effects.

**Fig 3.**
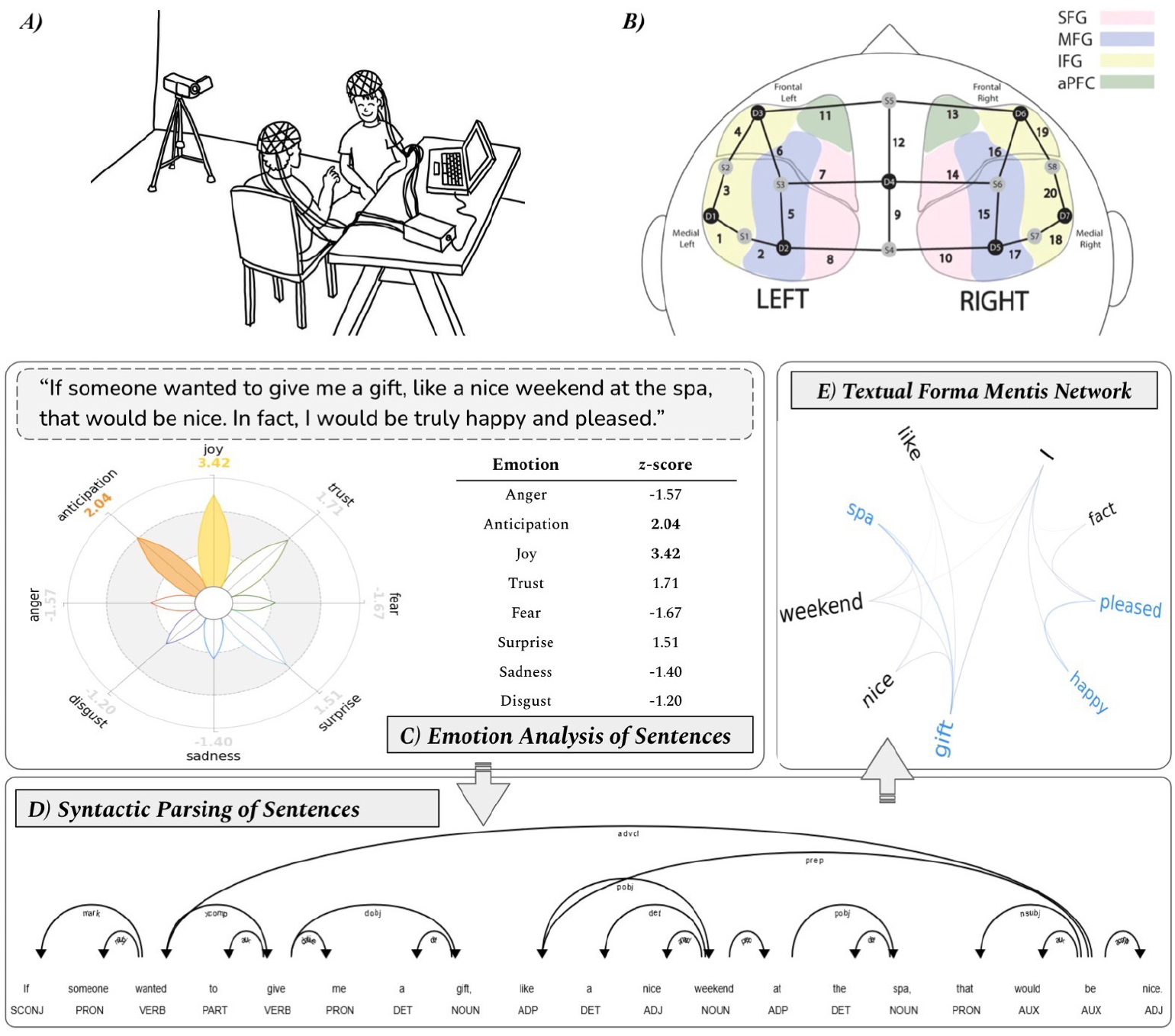
Schematic summary of the experimental procedure. ***A)*** Setup of devices and sitting arrangement of dyads during the experimental sessions. ***B)*** Image from [20]. Schematic diagram depicting the position of 20 functional near-infrared spectroscopy (fNIRS) channels and their corresponding positions to measure the activity of the superior frontal gyrus (SFG), middle frontal gyrus (MFG), inferior frontal gyrus (IFG), and anterior prefrontal cortex (aPFC). ***C)*** Emotional content of sentences computed using EmoAtlas [52]. ***D)*** Syntactic parsing of sentences. ***E)*** Representation of the syntactic/semantic structure of sentences using *textual forma mentis* networks [37].

### 5.4 Acquisition of Neural Data

fNIRS hyperscanning was used in all experimental tasks to monitor participants’ brain activity. Participants’ caps were set up with 8 LED sources emitting light at wavelengths of 760nm and 850nm, and 7 detectors, arranged following a standard prefrontal cortex montage [20, 79]. The overall configuration of sources and detectors resulted in 20 fNIRS channels for each participant, to monitor the activity of the prefrontal cortex. fNIRS channels were aggregated into the following regions of interest: anterior prefrontal cortex, superior frontal gyrus, left middle frontal gyrus, right middle frontal gyrus, left inferior frontal gyrus, and right inferior frontal gyrus (see Figure 3B). Due to their medial location within the prefrontal cortex and the spatial resolution limitations of fNIRS, the anterior prefrontal cortex and superior frontal gyrus were not divided into left and right portions, as was done for the other regions of interest. The placement of channels followed the standard international 10-20 electroencephalography layout [80]. Optode stabilizers were used to ensure that the distance between sources and detectors never exceeded 3 cm, to ensure a good signal-to-noise ratio [81]. For the data collection, a NIRSport2 device (NIRx Medical Technologies LLC) was used, with a sampling rate of 10.17 Hz.

### 5.5 Processing of Neural Data

Neural data were processed using *pyphysio* [82]. The quality of fNIRS signals was assessed using deep learning [83, 84]. More in detail, we used a convolutional neural network architecture trained to classify the quality of fNIRS segments. Motion artifacts were removed from fNIRS signals using spline interpolation [85] and wavelet filtering [86]. The processed data were subsequently converted to oxygenated (HbO) and deoxygenated hemoglobin (HbR) using the Beer-Lambert law [87]. We focused only on HbO values as in past studies (e.g., [19, 88]; see the Supplementary Materials for the results of the analyses on HbR values). Neural data from individual channels were ultimately aggregated into the regions of interest. The brain activity of each region of interest was obtained by computing the average of the normalized channels that composed each cluster [89]. To ensure good quality signals in the regions of interest, a region of interest signal was computed only if at least 2 channels with good quality were available [89].

### 5.6 Interpersonal Neural Synchrony

Interpersonal neural synchrony was computed between homologous brain regions of interest in the two conversationalists with Wavelet Transform Coherence (WTC) [19, 90, 91]. By using WTC, we assessed the coherence between individual fNIRS time series in each dyad and each region of interest as a function of frequency and time. WTC is particularly advantageous as it considers both phase-lagged correlations and in-phase correlations, enabling the assessment of global coherence patterns of brain activity [19].

For each dyad, we obtained eighteen values of WTC: one for each region of interest in each of the three experimental conditions (natural conversation, role-play, and role reversal). WTC was computed across the frequencies from 0.01 to 0.20 Hz in steps of 0.01 Hz [15, 92]. The final WTC value was calculated by averaging across the entire spectrum of frequencies, ensuring an unbiased approach as we had no prior hypothesis about specific frequency bands.

Moreover, for each region of interest in each experimental condition, we computed WTC between surrogate dyads, i.e., randomly paired participants taken from different real dyads. In this way, we obtain a “control” value of WTC derived from participants involved in a social task but that did not interact with one another directly. To ensure comparability, we performed a single permutation of the dyads, resulting in the same number of data points as the true dyads (i.e., 42 surrogate dyads). It is worth noting that these surrogate dyads are artificial and not experimentally reproduced or accessed in the lab. Surrogate dyads come from combining signals from individuals coming from different pairs tested in the lab. Thus, surrogate dyads have experimental conditions, regions of interest, and WTC scores but they do not have dialogues and, hence, no network or emotional feature.

### 5.7 Emotional Content of Dialogues

To analyze the emotional content in the transcribed dialogues, we employed EmoAtlas [52], a framework building TFMNs and performing emotional profiling based on Plutchick’s theory [40] (see Figure 3C). EmoAtlas operationalizes the quantification of eight emotions in text: anger, anticipation, disgust, fear, joy, sadness, surprise, and trust. For each emotion, the tool provides a *z* -score, indicating the intensity of that specific emotion within the dialogue compared to random assemblies of words from the underlying psychological data. Consider a text containing *m* emotional words. Random assemblies are created by sampling uniformly at random *m* words from the emotional lexicon of EmoAtlas. Because of uniform sampling, random assemblies will naturally reflect frequency effects in the emotional lexicon (e.g. there are more words eliciting trust rather than disgust). The library repeats random sampling 1000 times, creating a distribution of emotional word counts 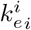 of finding that on the i-th sampling, 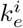 random words elicited the emotion *e*. Emotional *z* -scores are then computed as the observed counts of *m*_*e*_ words eliciting emotion *e* among the *m* ones found in text:

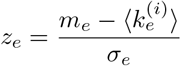

where *σ* is the error margin attributed to the sample mean computed over iterations is.

EmoAtlas was chosen due to its similar or superior performance compared to state-of-the-art natural language processing techniques [52].

To represent the emotional content of the dialogue at the dyad-level, we aggregated the *z* -scores for each emotion across participants within the (social) dyad.

In some cases, EmoAtlas is unable to assign reliable *z* -scores due to the low number of words in the provided text. In these cases (*n* = 2), emotional *z* -scores were noted as missing data.

### 5.8 Syntactic/Semantic Structure of Dialogues

To analyze the syntactic/semantic structure of dialogues, we employed *textual forma mentis* (Latin for “mindset”) networks (TFMNs) [37], built via EmoAtlas [52] (see Figure 3D and 3E). TFMNs map the associative knowledge of dialogues using network theory principles, wherein words/concepts serve as nodes associated by syntactic (specifications) or semantic (synonyms) links. TFMNs split any text into sentences and, then, sentence by sentence, they link words if they are nearby (at distance *K* ≤ 4) on the syntactic parsing tree of the sentence itself. TFMNs can, thus, spot specifications between words that are not adjacent or nearby in texts (i.e., separated by a few other words). This crucial difference makes TFMNs better at capturing syntactic relationships compared to word co-occurrence networks [93]. The syntactic parsing tree [37] (i.e., a tree graph determining syntactic dependencies) is computed sentence by sentence through a pre-trained AI model (from spaCy, see [52]). The *K* threshold is needed to link only syntactically close concepts and can be tuned by the experimenter. The value *K* ≤ 4 was selected in agreement with past findings showing in English syntactic distances between words mostly around 3 because of language optimization effects [94]. The syntactic network extracted by linking non-stopwords (at distance *K leq*4 on the parsing tree) is then enriched with synonyms and psychological emotional data to become a multiplex feature-rich network, where links can be either syntactic or semantic and words can be labeled as positive/negative/neutral and as eliciting one or more emotions (see [37, 50, 52]).

After computing the semantic network for each dyad in every condition, we extracted the following network measures [35, 95]: the total number of nodes |*V* | , the total number of edges| *E* |, the average local clustering, the number of connected components |*C* | , and degree assortativity (see Figure 4.). The total number of nodes| *V* |and |*E* |edges correspond to the number of words and syntactic/semantic associations included within the network, respectively. For instance, in a network representing a text corpus, |*V* | would represent the count of distinct words or concepts, while |*E* |would indicate how these words are connected based on their syntactic or semantic relationships. Average local clustering is an index of the likelihood that the neighbors of a node are linked with each other. Thus, average clustering measures the tendency of nodes to form clusters. In a semantic network, higher average clustering indicates that words or concepts tend to be densely interconnected within localized groups. The number of connected components |*C* | indicates the count of subsets within the network where there is a path between every pair of nodes. In other words, if the network consists of three groups of words where each group is internally connected but has no connections with the other groups, then the number of |*C* | would be 3. Finally, degree assortativity measures the association among nodes of similar weight, where weight can represent the number of connections (degree) a node has in the network. Higher degree assortativity values signify that nodes with a high degree (i.e., more connections) are linked to nodes with similarly high degrees.

**Fig 4.**
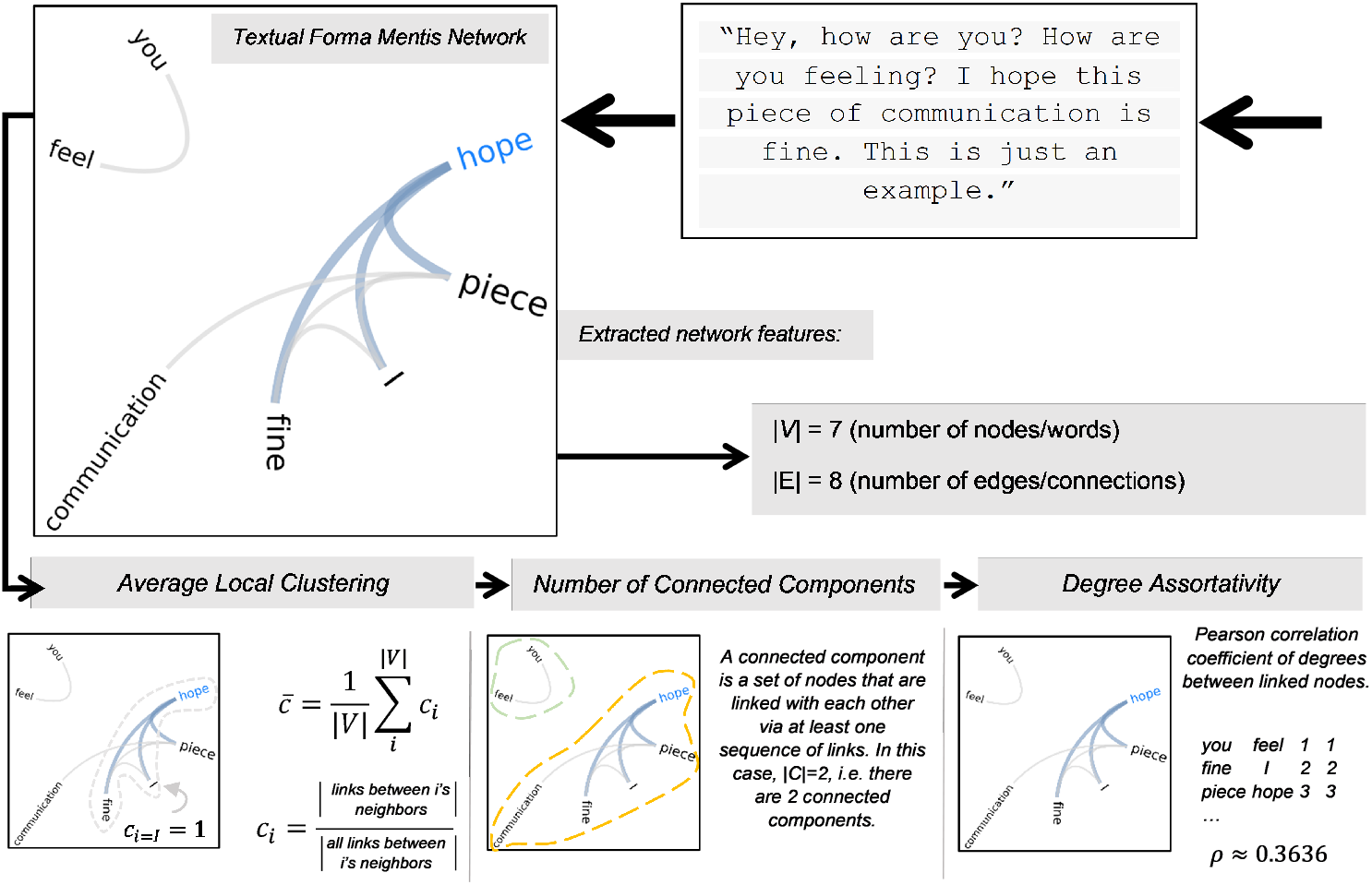
Network measures extracted from the semantic network built using *textual forma mentis* networks (TFMNs). From the semantic network, we extracted the total number of nodes |*V*|, the total number of edges|*E*|, the average local clustering, the number of connected components |*C*|, and degree assortativity.

For all syntactic/semantic metrics, individual values were aggregated to represent dyad-level data in each experimental condition.

### 5.9 Data Analysis

The analytical plan was divided into four phases:

1. A preliminary analysis of interpersonal neural synchrony across regions of interest;
2. An analysis of how the emotional content of dialogues relates to interpersonal neural synchrony;
3. An analysis of how the syntactic/semantic structure of dialogues relates to interpersonal neural synchrony;
4. An analysis of how the emotional content and the syntactic/semantic structure of dialogues together relate to interpersonal neural synchrony.

Initially, we compared the WTC in specific regions of interest between true *versus* surrogate dyads. To do so, six linear mixed models [96] with WTC scores as the dependent variable, dyad type (true *versus* surrogate) as fixed effect, and experimental condition and dyad ID as random effects were computed, one for each region of interest. After the statistical tests, all regions of interest with a significant difference in WTC scores between real and surrogate data were deemed eligible for the subsequent statistical analyses.

To investigate the association between emotional *z* -scores and interpersonal neural synchrony, we conducted a series of linear mixed models. In the first regression, we predicted WTC scores in one brain region of interest at a time. In this phase, for each region of interest with above-chance levels of synchrony, we used WTC scores as the dependent variable, the eight aggregated emotion *z* -scores as fixed effects, and experimental condition and dyad ID as random effects.

The same analysis was repeated by using the whole dataset to predict WTC scores across all subregions of the prefrontal cortex using the emotion *z* -scores. In this linear mixed model, we used WTC scores as the dependent variable, the eight aggregated emotion *z* -scores as fixed effects, and the region of interest, experimental condition, and dyad ID as random effects.

The same analytical approach was used to investigate the association between syntactic/semantic structure of dialogues and interpersonal neural synchrony. In this case, the TFMNs measures were used as fixed predictors of WTC in individual regions of interest and, subsequently, in the whole prefrontal cortex.

Finally, we combined the emotion *z* -scores and TFMNs’ syntactic/semantic structural features to predict interpersonal neural synchrony in the whole prefrontal cortex.

In this analysis, we used a linear mixed model with WTC scores as the dependent variable, emotion *z* -scores and syntactic/semantic metrics as fixed effects, and the region of interest, experimental condition, and dyad ID as random effects.

Before conducting the linear mixed models, we mean-centered the continuous fixed predictors of interest. In the analysis, to control for Type I errors due to multiple comparisons, we adopted Benjamini-Hochberg false discovery rate correction.

## Acknowledgments

Not applicable.

## Declarations

### Ethics approval

Data collections were approved by the University of Trento (2022-059) and by Nanyang Technological University (NTU-IRB-2021-03-013).

### Consent to participate

The conduction of the experiment followed the guidelines provided by the Declaration of Helsinki and informed consent was obtained from all participants.

### Data availability

Data and materials are available upon request to the corresponding author.

### Authors’ contributions

Conceptualization: AC, MS, GE; Methodology: AC, MS, AB; Formal analysis: AC, MS; Investigation: AC, ML; Data Curation: AC, ML; Writing – Original Draft: AC, MS; Writing – Review & Editing: AC, MS, ML, AB, GE; Supervision: GE.

## References

[1] Bowlby, J. & Holmes, J. A Secure Base (Routledge, London, 2012).

[2] Ferjan Ramírez, N., Lytle, S. R. & Kuhl, P. K. Parent coaching increases conversational turns and advances infant language development. Proceedings of the National Academy of Sciences 117, 3484–3491 (2020).

[3] Saint-Georges, C. et al. Motherese in interaction: at the cross-road of emotion and cognition?(a systematic review). PloS one 8, e78103 (2013).

[4] Johnson, M. H., Dziurawiec, S., Ellis, H. & Morton, J. Newborns’ preferential tracking of face-like stimuli and its subsequent decline. Cognition 40, 1–19 (1991).

[5] Bolis, D., Dumas, G. & Schilbach, L. Interpersonal attunement in social interactions: from collective psychophysiology to inter-personalized psychiatry and beyond. Philosophical Transactions of the Royal Society B 378, 20210365 (2023).

[6] Holt-Lunstad, J., Smith, T. B. & Layton, J. B. Social relationships and mortality risk: a meta-analytic review. PLoS medicine 7, e1000316 (2010).

[7] Bojczyk, K. E., Davis, A. E. & Rana, V. Mother–child interaction quality in shared book reading: Relation to child vocabulary and readiness to read. Early childhood research quarterly 36, 404–414 (2016).

[8] Atzil, S. & Gendron, M. Bio-behavioral synchrony promotes the development of conceptualized emotions. Current opinion in psychology 17, 162–169 (2017).

[9] Feldman, R. Parent–infant synchrony and the construction of shared timing; physiological precursors, developmental outcomes, and risk conditions. Journal of Child psychology and Psychiatry 48, 329–354 (2007).

[10] Feldman, R. Bio-behavioral synchrony: A model for integrating biological and microsocial behavioral processes in the study of parenting. Parenting 12, 154–164 (2012).

[11] Feldman, R. The neurobiology of human attachments. Trends in cognitive sciences 21, 80–99 (2017).

[12] Carollo, A., Lim, M., Aryadoust, V. & Esposito, G. Interpersonal synchrony in the context of caregiver-child interactions: A document co-citation analysis. Frontiers in psychology 12, 701824 (2021).

[13] Montague, P. R. et al. Hyperscanning: simultaneous fmri during linked social interactions (2002).

[14] Astolfi, L. et al. Neuroelectrical hyperscanning measures simultaneous brain activity in humans. Brain topography 23, 243–256 (2010).

[15] Lim, M., Carollo, A., Bizzego, A., Chen, A. S. & Esposito, G. Culture, sex and social context influence brain-to-brain synchrony: an fnirs hyperscanning study. BMC psychology 12, 350 (2024).

[16] Morgan, J. K. et al. Mother–child neural synchronization is time linked to mother–child positive affective state matching. Social Cognitive and Affective Neuroscience 18, nsad001 (2023).

[17] Nguyen, T., Kungl, M. T., Hoehl, S., White, L. O. & Vrtička, P. Visualizing the invisible tie: Linking parent–child neural synchrony to parents’ and children’s attachment representations. Developmental Science e13504 (2023).

[18] Carollo, A. & Esposito, G. Hyperscanning literature after two decades of neuroscientific research: a scientometric review. Neuroscience 551, 345–354 (2024).

[19] Nguyen, T. et al. The effects of interaction quality on neural synchrony during mother-child problem solving. cortex 124, 235–249 (2020).

[20] Azhari, A. et al. Parenting stress undermines mother-child brain-to-brain synchrony: A hyperscanning study. Scientific reports 9, 1–9 (2019).

[21] Azhari, A., Bizzego, A. & Esposito, G. Parent–child dyads with greater parenting stress exhibit less synchrony in posterior areas and more synchrony in frontal areas of the prefrontal cortex during shared play. Social Neuroscience 17, 520–531 (2022).

[22] Balconi, M. & Vanutelli, M. E. Cooperation and competition with hyperscanning methods: review and future application to emotion domain. Frontiers in computational neuroscience 11, 86 (2017).

[23] Cornejo, C., Cuadros, Z., Morales, R. & Paredes, J. Interpersonal coordination: methods, achievements, and challenges. Frontiers in psychology 8, 1685 (2017).

[24] Czeszumski, A. et al. Hyperscanning: a valid method to study neural inter-brain underpinnings of social interaction. Frontiers in Human Neuroscience 14, 39 (2020).

[25] Lopes, P. N., Salovey, P., Côté, S., Beers, M. & Petty, R. E. Emotion regulation abilities and the quality of social interaction. Emotion 5, 113 (2005).

[26] Twenge, J. M., Baumeister, R. F., DeWall, C. N., Ciarocco, N. J. & Bartels, J. M. Social exclusion decreases prosocial behavior. Journal of personality and social psychology 92, 56 (2007).

[27] Russell, J. A. A circumplex model of affect. Journal of personality and social psychology 39, 1161 (1980).

[28] Nummenmaa, L. et al. Emotions promote social interaction by synchronizing brain activity across individuals. Proceedings of the National Academy of Sciences 109, 9599–9604 (2012).

[29] Santamaria, L. et al. Emotional valence modulates the topology of the parent-infant inter-brain network. NeuroImage 207, 116341 (2020).

[30] Zhu, L. et al. Neural mechanisms of social emotion perception: an eeg hyper-scanning study, 199–206 (IEEE, 2018).

[31] Hoehl, S., Fairhurst, M. & Schirmer, A. Interactional synchrony: signals, mechanisms and benefits. Social cognitive and affective neuroscience 16, 5–18 (2021).

[32] Aitchison, J. Words in the mind: An introduction to the mental lexicon (John Wiley & Sons, 2012).

[33] Chan, K. Y. & Vitevitch, M. S. Network structure influences speech production. Cognitive science 34, 685–697 (2010).

[34] Vitevitch, M. S., Pisoni, D. B., Soehlke, L. & Foster, T. A. Using complex networks in the hearing sciences. Ear and Hearing 45, 1–9 (2024).

[35] Stella, M. Cognitive network science for understanding online social cognitions: A brief review. Topics in Cognitive Science 14, 143–162 (2022).

[36] Tausczik, Y. R. & Pennebaker, J. W. The psychological meaning of words: Liwc and computerized text analysis methods. Journal of language and social psychology 29, 24–54 (2010).

[37] Stella, M., De Nigris, S., Aloric, A. & Siew, C. S. Forma mentis networks quantify crucial differences in stem perception between students and experts. PloS one 14, e0222870 (2019).

[38] Ferrer-i Cancho, R., Gómez-Rodríguez, C., Esteban, J. L. & Alemany-Puig, L. Optimality of syntactic dependency distances. Physical Review E 105, 014308 (2022).

[39] Polkinghorne, D. E. Language and meaning: Data collection in qualitative research. Journal of counseling psychology 52, 137 (2005).

[40] Plutchik, R. in A general psychoevolutionary theory of emotion 3–33 (Elsevier, 1980).

[41] Mohammad, S. M. in Sentiment analysis: Detecting valence, emotions, and other affectual states from text 201–237 (Elsevier, 2016).

[42] Castro, N., Stella, M. & Siew, C. S. Quantifying the interplay of semantics and phonology during failures of word retrieval by people with aphasia using a multiplex lexical network. Cognitive science 44, e12881 (2020).

[43] Ciampelli, S. et al. Syntactic network analysis in schizophrenia-spectrum disorders. Schizophrenia Bulletin 49, S172–S182 (2023).

[44] Stella, M. et al. Cognitive modelling of concepts in the mental lexicon with multilayer networks: Insights, advancements, and future challenges. Psychonomic Bulletin & Review 1–24 (2024).

[45] Nguyen, T. et al. Neural synchrony in mother–child conversation: Exploring the role of conversation patterns. Social Cognitive and Affective Neuroscience 16, 93–102 (2021).

[46] Nguyen, T., Zimmer, L. & Hoehl, S. Your turn, my turn. neural synchrony in mother–infant proto-conversation. Philosophical Transactions of the Royal Society B 378, 20210488 (2023).

[47] Lim, M., Carollo, A., Bizzego, A., Chen, S. A. & Esposito, G. Decreased activation in left prefrontal cortex during role-play: an fnirs study of the psychodrama sociocognitive model. The Arts in Psychotherapy 102098 (2023).

[48] Huth, A. G., De Heer, W. A., Griffiths, T. L., Theunissen, F. E. & Gallant, J. L. Natural speech reveals the semantic maps that tile human cerebral cortex. Nature 532, 453–458 (2016).

[49] Yang, Y. et al. Unraveling lexical semantics in the brain: Comparing internal, external, and hybrid language models. Human Brain Mapping 45, e26546 (2024).

[50] Semeraro, A., Vilella, S., Ruffo, G. & Stella, M. Emotional profiling and cognitive networks unravel how mainstream and alternative press framed astrazeneca, pfizer and covid-19 vaccination campaigns. Scientific reports 12, 14445 (2022).

[51] Lim, M., Carollo, A., Bizzego, A., Chen, A. S. & Esposito, G. Synchrony within, synchrony without: establishing the link between interpersonal behavioural and brain-to-brain synchrony during role-play. Royal Society Open Science 11, 240331 (2024).

[52] Semeraro, A. et al. Emoatlas: An emotional network analyser of texts merging psychological lexicons, artificial intelligence and network science. PsyArxiv (2024).

[53] Cui, X., Bryant, D. M. & Reiss, A. L. Nirs-based hyperscanning reveals increased interpersonal coherence in superior frontal cortex during cooperation. Neuroimage 59, 2430–2437 (2012).

[54] Dziobek, I. et al. Neuronal correlates of altered empathy and social cognition in borderline personality disorder. Neuroimage 57, 539–548 (2011).

[55] Pinti, P. et al. The role of anterior prefrontal cortex (area 10) in face-to-face deception measured with fnirs. Social Cognitive and Affective Neuroscience 16, 129–142 (2021).

[56] Christ, S. E., Van Essen, D. C., Watson, J. M., Brubaker, L. E. & McDermott, K. B. The contributions of prefrontal cortex and executive control to deception: evidence from activation likelihood estimate meta-analyses. Cerebral cortex 19, 1557–1566 (2009).

[57] Ding, X. P. et al. The neural correlates of identity faking and concealment: an fmri study. PLoS One 7, e48639 (2012).

[58] Langleben, D. D. et al. Brain activity during simulated deception: an event-related functional magnetic resonance study. Neuroimage 15, 727–732 (2002).

[59] Kinreich, S., Djalovski, A., Kraus, L., Louzoun, Y. & Feldman, R. Brain-to-brain synchrony during naturalistic social interactions. Scientific reports 7, 17060 (2017).

[60] Grecucci, A., Giorgetta, C., Bonini, N. & Sanfey, A. G. Reappraising social emotions: the role of inferior frontal gyrus, temporo-parietal junction and insula in interpersonal emotion regulation. Frontiers in human neuroscience 7, 523 (2013).

[61] Frank, D. et al. Emotion regulation: quantitative meta-analysis of functional activation and deactivation. Neuroscience & Biobehavioral Reviews 45, 202–211 (2014).

[62] Roy, M., Shohamy, D. & Wager, T. D. Ventromedial prefrontal-subcortical systems and the generation of affective meaning. Trends in cognitive sciences 16, 147–156 (2012).

[63] Waugh, C. E., Lemus, M. G. & Gotlib, I. H. The role of the medial frontal cortex in the maintenance of emotional states. Social Cognitive and Affective Neuroscience 9, 2001–2009 (2014).

[64] Waugh, C. E., Hamilton, J. P. & Gotlib, I. H. The neural temporal dynamics of the intensity of emotional experience. Neuroimage 49, 1699–1707 (2010).

[65] Zhang, B. et al. Altered spontaneous neural activity in the precuneus, middle and superior frontal gyri, and hippocampus in college students with subclinical depression. BMC psychiatry 21, 1–10 (2021).

[66] Tabei, K.-i. et al. Inferior frontal gyrus activation underlies the perception of emotions, while precuneus activation underlies the feeling of emotions during music listening. Behavioural neurology 2015 (2015).

[67] Newman, M. Networks (Oxford university press, 2018).

[68] Kotz, S. A., Cappa, S. F., von Cramon, D. Y. & Friederici, A. D. Modulation of the lexical–semantic network by auditory semantic priming: An event-related functional mri study. Neuroimage 17, 1761–1772 (2002).

[69] Lau, E. F., Phillips, C. & Poeppel, D. A cortical network for semantics:(de) constructing the n400. Nature reviews neuroscience 9, 920–933 (2008).

[70] Rissman, J., Eliassen, J. C. & Blumstein, S. E. An event-related fmri investigation of implicit semantic priming. Journal of cognitive neuroscience 15, 1160–1175 (2003).

[71] Laufer, I., Negishi, M., Lacadie, C. M., Papademetris, X. & Constable, R. T. Dissociation between the activity of the right middle frontal gyrus and the middle temporal gyrus in processing semantic priming. PloS one 6, e22368 (2011).

[72] Sachs, O. et al. How different types of conceptual relations modulate brain activation during semantic priming. Journal of Cognitive Neuroscience 23, 1263–1273 (2011).

[73] Raposo, A., Moss, H. E., Stamatakis, E. A. & Tyler, L. K. Repetition suppression and semantic enhancement: an investigation of the neural correlates of priming. Neuropsychologia 44, 2284–2295 (2006).

[74] Dixon, M. L., Thiruchselvam, R., Todd, R. & Christoff, K. Emotion and the prefrontal cortex: An integrative review. Psychological bulletin 143, 1033 (2017).

[75] Hertrich, I., Dietrich, S., Blum, C. & Ackermann, H. The role of the dorsolateral prefrontal cortex for speech and language processing. Frontiers in human neuroscience 15, 645209 (2021).

[76] Forbes, C. E. & Grafman, J. The role of the human prefrontal cortex in social cognition and moral judgment. Annual review of neuroscience 33, 299–324 (2010).

[77] Alonso-Martin, F., Malfaz, M., Sequeira, J., Gorostiza, J. F. & Salichs, M. A. A multimodal emotion detection system during human–robot interaction. Sensors 13, 15549–15581 (2013).

[78] Levinson, S. C. Action formation and ascription. The handbook of conversation analysis 101–130 (2012).

[79] Bizzego, A., Gabrieli, G., Azhari, A., Lim, M. & Esposito, G. Dataset of parent-child hyperscanning functional near-infrared spectroscopy recordings. Scientific Data 9, 625 (2022).

[80] Homan, R. W., Herman, J. & Purdy, P. Cerebral location of international 10–20 system electrode placement. Electroencephalography and clinical neurophysiology 66, 376–382 (1987).

[81] Pinti, P. et al. The present and future use of functional near-infrared spectroscopy (fnirs) for cognitive neuroscience. Annals of the New York Academy of Sciences 1464, 5–29 (2020).

[82] Bizzego, A., Battisti, A., Gabrieli, G., Esposito, G. & Furlanello, C. pyphysio: A physiological signal processing library for data science approaches in physiology. SoftwareX 10, 100287 (2019).

[83] Bizzego, A., Neoh, M., Gabrieli, G. & Esposito, G. A machine learning perspective on fnirs signal quality control approaches. IEEE Transactions on Neural Systems and Rehabilitation Engineering 30, 2292–2300 (2022).

[84] Bizzego, A., Gabrieli, G. & Esposito, G. Deep neural networks and transfer learning on a multivariate physiological signal dataset. Bioengineering 8, 35 (2021).

[85] Scholkmann, F., Spichtig, S., Muehlemann, T. & Wolf, M. How to detect and reduce movement artifacts in near-infrared imaging using moving standard deviation and spline interpolation. Physiological measurement 31, 649 (2010).

[86] Molavi, B. & Dumont, G. A. Wavelet-based motion artifact removal for functional near-infrared spectroscopy. Physiological measurement 33, 259 (2012).

[87] Delpy, D. & Cope, M. Quantification in tissue near–infrared spectroscopy. Philosophical Transactions of the Royal Society of London. Series B: Biological Sciences 352, 649–659 (1997).

[88] Reindl, V., Gerloff, C., Scharke, W. & Konrad, K. Brain-to-brain synchrony in parent-child dyads and the relationship with emotion regulation revealed by fnirs-based hyperscanning. NeuroImage 178, 493–502 (2018).

[89] Bizzego, A., Azhari, A. & Esposito, G. Reproducible inter-personal brain coupling measurements in hyperscanning settings with functional near infra-red spectroscopy. Neuroinformatics 20, 665–675 (2022).

[90] Chang, C. & Glover, G. H. Time–frequency dynamics of resting-state brain connectivity measured with fmri. Neuroimage 50, 81–98 (2010).

[91] Grinsted, A., Moore, J. C. & Jevrejeva, S. Application of the cross wavelet transform and wavelet coherence to geophysical time series. Nonlinear processes in geophysics 11, 561–566 (2004).

[92] Holper, L., Scholkmann, F. & Wolf, M. Between-brain connectivity during imitation measured by fnirs. Neuroimage 63, 212–222 (2012).

[93] Quispe, L. V., Tohalino, J. A. & Amancio, D. R. Using virtual edges to improve the discriminability of co-occurrence text networks. Physica A: Statistical Mechanics and its Applications 562, 125344 (2021).

[94] i Cancho, R. F., Solé, R. V. & Köhler, R. Patterns in syntactic dependency networks. Physical Review E 69, 051915 (2004).

[95] Siew, C. S., Wulff, D. U., Beckage, N. M. & Kenett, Y. N. Cognitive network science: A review of research on cognition through the lens of network representations, processes, and dynamics. Complexity (2019).

[96] Brown, V. A. An introduction to linear mixed-effects modeling in r. Advances in Methods and Practices in Psychological Science 4, 2515245920960351 (2021).

