## Supplemental Tables 1, 2, 3 for "Emotional Content and Semantic Structure of Dialogues Are Associated With Interpersonal Neural Synchrony in the Prefrontal Cortex"

In the Supplementary Materials, we repeated the statistical analyses conducted in the main manuscript, focusing specifically on the values for the deoxygenated hemoglobin component (HbR).

### 1. Interpersonal Neural Synchrony Across True and Surrogate Dyads

As was done for the HbO values in the manuscript, we conducted the same statistical analyses on the HbR values. Specifically, to identify regions of interest that exhibited significant differences between true and surrogate dyads, we performed six linear mixed-effects models – one for each individual region of interest. In these models, WTC scores was used as the dependent variable, dyad type (true versus surrogate) was included as a fixed effect, while experimental condition and dyad ID were treated as random effects. None of the models showed statistically significant differences in WTC values between dyad types ($q > .05$).

### 2. Emotional Content and Interpersonal Neural Synchrony in Individual Regions of Interest

To examine the relationship between the emotional content of dialogues and interpersonal neural synchrony, we performed three separate linear mixed-effects models, analyzing data from the superior frontal gyrus, the left middle frontal gyrus, and the right middle frontal gyrus individually. In each model, WTC scores based on HbR values were used as the dependent variable, emotional $z$-scores were included as fixed effects, and experimental condition and dyad ID were treated as random effects. None of the models revealed a statistically significant effect of emotional $z$-scores on WTC values ($q > .05$; see Table S1 for an overview of the results).

### 3. Syntactic/Semantic Structure of Dialogues and Interpersonal Neural Synchrony in Individual Regions of Interest

To examine the relationship between the syntactic/semantic structure of dialogues and interpersonal neural synchrony, we performed three separate linear mixed-effects models, analyzing data from the superior frontal gyrus, the left middle frontal gyrus, and the right middle frontal gyrus individually. In each model, WTC scores based on HbR values were used as the dependent variable, syntactic/semantic metrics were included as fixed

| Predictor | Estimate ($\beta$) | Standard Error | $t$-value | $p$-value | $q$-value |
|---|---|---|---|---|---|
| **Interpersonal Neural Synchrony on the Superior Frontal Gyrus** | | | | | |
| **Anger** | 0.0027 | 0.008 | 0.340 | .734 | .980 |
| **Anticipation** | -0.0073 | 0.007 | -1.083 | .281 | .807 |
| **Disgust** | -0.0025 | 0.008 | -0.323 | .747 | .960 |
| **Fear** | -0.0061 | 0.007 | -0.885 | .378 | .807 |
| **Joy** | 0.0008 | 0.006 | 0.136 | .892 | .980 |
| **Sadness** | 0.0138 | 0.007 | 2.117 | .037 | .247 |
| **Surprise** | 0.0103 | 0.007 | 1.518 | .132 | .594 |
| **Trust** | -0.0005 | 0.006 | -0.078 | .938 | .980 |
| **Interpersonal Neural Synchrony on the Left Middle Frontal Gyrus** | | | | | |
| **Anger** | 0.0018 | 0.008 | 0.228 | .820 | .980 |
| **Anticipation** | 0.0078 | 0.007 | 1.159 | .249 | .807 |
| **Disgust** | 0.0014 | 0.008 | 0.180 | .858 | .901 |
| **Fear** | 0.0041 | 0.007 | 0.612 | .542 | .921 |
| **Joy** | 0.0050 | 0.006 | 0.813 | .418 | .807 |
| **Sadness** | 0.0033 | 0.007 | 0.505 | .614 | .921 |
| **Surprise** | -0.0002 | 0.007 | -0.027 | .979 | .980 |
| **Trust** | 0.0026 | 0.006 | 0.430 | .668 | .921 |
| **Interpersonal Neural Synchrony on the Right Middle Frontal Gyrus** | | | | | |
| **Anger** | 0.0002 | 0.008 | 0.025 | .980 | .980 |
| **Anticipation** | 0.0091 | 0.007 | 1.270 | .207 | .798 |
| **Disgust** | -0.0064 | 0.008 | -0.814 | .417 | .807 |
| **Fear** | 0.0060 | 0.007 | 0.848 | .398 | .807 |
| **Joy** | -0.0065 | 0.006 | -1.024 | .308 | .807 |
| **Sadness** | 0.0127 | 0.007 | 1.889 | .062 | .333 |
| **Surprise** | -0.0043 | 0.007 | -0.601 | .549 | .921 |
| **Trust** | -0.0033 | 0.006 | -0.528 | .598 | .921 |

Table 1: Summary of fixed effects related to the emotion $z$-scores in the linear mixed models conducted to predict interpersonal neural synchrony in individual regions of interest (i.e., superior frontal gyrus, left middle frontal gyrus, right middle frontal gyrus). For each emotion, we reported the estimate, the standard error, the $t$-value, the $p$-value, and the $q$-value. Statistically significant $q$-values are noted in bold.

effects, and experimental condition and dyad ID were treated as random effects.

One out of the three models showed a significant effect of syntactic/semantic properties of dialogues on interpersonal neural synchrony (see Table S2 for a summary of the results). Specifically, the model on data from the superior middle frontal gyrus showed a significant effect of the degree assortativity ($\beta = -0.019$, $SE$ = 0.006, $t$(112.90) = -2.917, $p$ = .004, $q$ = .016) on WTC scores. The $AIC$ of the model equaled to -291.95 and the $R^2_{marginal}$ was equal to 12.03%. In contrast, syntactic/semantic metrics were not significant predictors of interpersonal neural synchrony in the models on data from the left and right middle frontal gyri ($q > .05$).

| Predictor | Estimate ($\beta$) | Standard Error | $t$-value | $p$-value | $q$-value |
|---|---|---|---|---|---|
| **Interpersonal Neural Synchrony on the Superior Frontal Gyrus** | | | | | |
| **Nodes** | 0.0107 | 0.005 | 1.974 | .051 | .153 |
| **Average Local Clustering** | -0.0066 | 0.005 | -1.222 | .224 | .428 |
| **Connected Components** | -0.0010 | 0.006 | -0.152 | .879 | .902 |
| **Degree Assortativity** | -0.0189 | 0.006 | -2.917 | .004 | **.016** * |
| **Interpersonal Neural Synchrony on the Left Middle Frontal Gyrus** | | | | | |
| **Nodes** | 0.0055 | 0.006 | 0.959 | .340 | .526 |
| **Average Local Clustering** | 0.0007 | 0.006 | 0.124 | .902 | .902 |
| **Connected Components** | 0.0052 | 0.007 | 0.768 | .444 | .605 |
| **Degree Assortativity** | -0.0063 | 0.007 | -0.935 | .352 | .528 |
| **Interpersonal Neural Synchrony on the Right Middle Frontal Gyrus** | | | | | |
| **Nodes** | -0.0072 | 0.006 | -1.198 | .233 | .438 |
| **Average Local Clustering** | -0.0075 | 0.0059 | -1.277 | .204 | .438 |
| **Connected Components** | 0.0021 | 0.007 | 0.295 | .768 | .902 |
| **Degree Assortativity** | 0.0013 | 0.007 | 0.182 | .856 | .902 |

Table 2: Summary of fixed effects related to the syntactic/semantic properties of dialogues in the linear mixed models conducted to predict interpersonal neural synchrony in individual regions of interest (i.e., superior frontal gyrus, left middle frontal gyrus, right middle frontal gyrus). For each syntactic/semantic metric, we reported the estimate, the standard error, the $t$-value, the $p$-value, and the $q$-value. Statistically significant $q$-values are noted in bold. (* $q < .05$).

### 4. Analyses on the Whole Prefrontal Cortex

The analyses were also performed using data from all three regions of interest combined. We conducted three linear mixed-effects models, with WTC scores as the dependent variable and region of interest, experimental condition, and dyad ID as random effects. In the first model, emotional $z$-scores were included as the only fixed effects. In the second model, syntactic/semantic metrics were included as the only fixed effects. In the third model, both emotional and syntactic/semantic information were included as fixed effects. None of the three models revealed statistically significant effects of the emotional or syntactic/semantic properties of the dialogues on interpersonal neural synchrony ($q > .05$; see Table S3 for an overview of the results).

| **Predictor** | **Estimate ($\beta$)** | **Standard Error** | ***t*-value** | ***p*-value** | ***q*-value** |
|---|---|---|---|---|---|
| | | **Emotional *z*-Scores** | | | |
| **Anger** | 0.0009 | 0.005 | 0.186 | .852 | .933 |
| **Anticipation** | 0.0033 | 0.004 | 0.838 | .403 | .837 |
| **Disgust** | -0.0020 | 0.004 | -0.452 | .651 | .933 |
| **Fear** | 0.0013 | 0.004 | 0.329 | .743 | .933 |
| **Joy** | -0.0003 | 0.004 | -0.070 | .944 | .944 |
| **Sadness** | 0.0102 | 0.004 | 2.664 | .008 | .057 |
| **Surprise** | 0.0020 | 0.004 | 0.506 | .613 | .933 |
| **Trust** | -0.0004 | 0.004 | -0.122 | .903 | .937 |
| | | **Syntactic/Semantic Properties** | | | |
| **Nodes** | 0.0032 | 0.003 | 0.931 | .353 | .795 |
| **Average Local Clustering** | -0.0044 | 0.003 | -1.295 | .197 | .653 |
| **Connected Components** | 0.0023 | 0.004 | 0.579 | .564 | .933 |
| **Degree Assortativity** | -0.0081 | 0.004 | -2.008 | .050 | .207 |
| | | **Emotional *z*-Scores and Syntactic/Semantic Properties** | | | |
| **Anger** | 0.0015 | 0.005 | 0.311 | .756 | .933 |
| **Anticipation** | 0.0039 | 0.004 | 0.956 | .340 | .795 |
| **Disgust** | -0.0017 | 0.005 | -0.381 | .704 | .933 |
| **Fear** | 0.0022 | 0.004 | 0.539 | .591 | .933 |
| **Joy** | -0.0006 | 0.004 | -0.172 | .864 | .933 |
| **Sadness** | 0.0095 | 0.004 | 2.331 | .021 | .112 |
| **Surprise** | 0.0013 | 0.0040 | 0.329 | .743 | .933 |
| **Trust** | -0.0008 | 0.004 | -0.222 | .824 | .933 |
| **Nodes** | 0.0034 | 0.003 | 1.014 | .312 | .795 |
| **Average Local Clustering** | -0.0053 | 0.003 | -1.554 | .122 | .470 |
| **Connected Components** | -0.0010 | 0.004 | -0.238 | .812 | .933 |
| **Degree Assortativity** | -0.0052 | 0.004 | -1.237 | .218 | .652 |

Table 3: Summary of fixed effects in the linear mixed models conducted to predict interpersonal neural synchrony using data from all regions of interest (i.e., superior frontal gyrus, left middle frontal gyrus, right middle frontal gyrus). For each predictor, we reported the estimate, the standard error, the $t$-value, the $p$-value, and the $q$-value. Statistically significant $q$-values are noted in bold.
